# Applying a gene-suite approach to examine the physiological status of wild-caught walleye (*Sander vitreus*)

**DOI:** 10.1101/2020.02.29.971374

**Authors:** Jennifer D. Jeffrey, Hunter Carlson, Dale Wrubleski, Eva C. Enders, Jason R. Treberg, Ken M. Jeffries

## Abstract

Molecular techniques have been increasingly used in a conservation physiology framework to provide valuable information regarding the mechanisms underlying responses of wild organisms to environmental and anthropogenic stressors. In the present study, we developed a reference gill transcriptome for walleye (*Sander vitreus*) allowing us to pair a gene-suite approach with multivariate statistics to examine the physiological status of wild-caught walleye. For molecular analyses of wild fish, the gill is a useful target for conservation studies, not only because of its importance as an indicator of the physiological status of fish but also because it can be biopsied non-lethally. Walleye were non-lethally sampled following short- (∼1.5 month) and long-term (∼3.5 month) holding in the Delta Marsh, that is located south of Lake Manitoba in Manitoba, Canada. Large-bodied walleye are held in the Delta Marsh from late April to early August by exclusion screens used to protect the marsh from invasive common carp (*Cyprinus carpio*), exposing fish to potentially stressful water quality conditions. Principal components analysis (PCA) revealed patterns of transcript abundance consistent with exposure of fish to increasingly high temperature and low oxygen conditions with longer holding in the marsh. For example, longer-term holding in the marsh was associated with increases in the mRNA levels of heat shock proteins and a shift in the mRNA abundance of aerobic to anaerobic metabolic genes. Overall, the results of the present study suggest that walleye held in the Delta Marsh may be exhibiting sub-lethal responses to high temperature and low oxygen conditions and provides valuable information for managers invested in mediating these impacts to a local species of conservation concern. More broadly, we highlight the usefulness of pairing transcriptomic techniques with multivariate statistics to address potential confounding factors that can affect measured physiological responses of wild-caught fish.

**Lay summary:** Non-lethal molecular techniques were used to assess the physiological status of wild-caught walleye confined in the Delta Marsh, Manitoba, Canada, because of common carp exclusion screens. Walleye sampled during the warmest months of the year exhibited responses to elevated temperature and low oxygen conditions, suggesting sub-lethal effects of local conditions.

## Introduction

Conservation physiology has resulted in the increased use of physiological tools to assess the effects of environmental change and anthropogenic stressors on organismal responses (Wikelski and Cooke, 2006; Cooke *et al*., 2013). Incorporating physiological knowledge with ecological modeling can provide valuable information regarding the mechanisms underlying conservation problems (Cooke *et al*., 2013), as well as short-term indicators of organismal stress that may not be evident by long-term population and community metrics (Jeffrey *et al*., 2015). However, studying the physiological status of wild organisms can be challenging, with the potential for confounding factors induced by capture and handling stressors (Fig. 1). For wild fishes, capture techniques, such as gill or trammel netting, trap netting, electrofishing, and angling, may require that fish remain in nets or live wells for extended periods of time prior to sampling. Thus, techniques used in the wild to capture fish could potentially affect various physiological responses, making it difficult to distinguish between responses to handling and capture stressors *versus* responses to the environment prior to capture. For instance, commonly used indicators of stress, such as plasma cortisol, begin to rise within minutes following exposure to a stressor and can rise to peak levels within 30–60 min in some fishes (Barton, 2002). Thus, physiological indicators that are sensitive to acute handling or capture stressors may not be representative of the environmental conditions experienced by a fish prior to capture. Furthermore, some severe acute stressors may mask more subtle physiological responses from exposure to chronic environmental stressors. This difficulty in distinguishing the physiological responses due to the environmental conditions prior to capture from those induced by capture and handling techniques is a challenge facing conservation physiologists.

**Figure 1.**
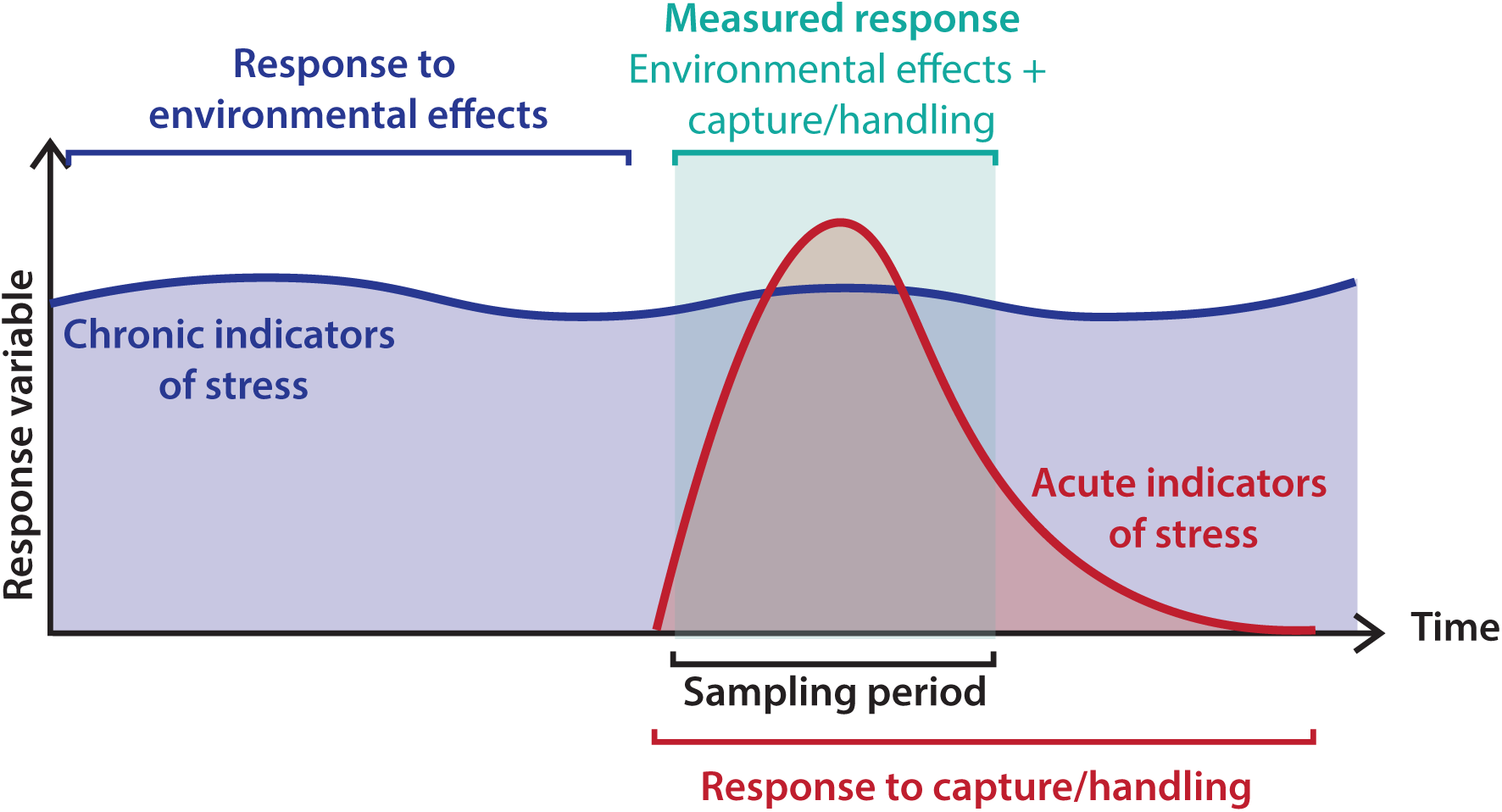
Visual representation of acute *versus* chronic indicators of stress. Acute indicators of stress can be sensitive to handling or capture stressors in wild fish. To best reflect the responses of wild fish to environmental stressors or conditions, indicators of chronic stress, that are less likely to be sensitive to handling or capture stressors, should be targeted.

Transcriptomic approaches have been increasingly used with wild fishes in the framework of conservation physiology (Connon *et al*., 2018). As the initial step in gene expression, gene transcription is a key regulator of physiological status and may provide insights into phenotypic plasticity and responses to environmental factors in wild fish (Alvarez *et al*., 2015; Oomen and Hutchings, 2017), constituting an important tool in the conservation physiology toolbox (Madliger *et al*., 2018). The increased affordability of RNA-sequencing (RNA-seq) technology has facilitated the development of *de novo* reference transcriptomes for non-model species making it possible to design whole-transcriptomic, or targeted gene-suite (i.e., with tens to hundreds of genes) studies to examine the physiological status of wild fish. For example, high-throughput qPCR approaches have been developed for characterizing pathogen infections and immune responses in wild Pacific salmon in British Columbia, Canada (e.g., Jeffries *et al*., 2014; Miller *et al*., 2014; Thakur *et al*., 2018; Tucker *et al*., 2018). Gene-suite studies can be preferable to whole-transcriptomic studies to reduce cost, increase sample sizes, simplify data analyses, and provide a more targeted approach that can translate easier to non-experts.

Walleye (*Sander vitreus*) is a species that could benefit from the increased availability of molecular tools and stress biomarkers due to their commercial value and as a species of concern in some locations throughout their distribution (Zuiden and Sharma, 2016; Government of Canada and Manitoba Government, 2018; Rypel *et al*., 2018). Walleye are a popular cool-water fish for recreational and commercial fishing that is widely distributed in North America. North America’s second largest commercial inland fishery is located in the province of Manitoba, of which walleye make up an important component, highlighting their local economic value (Sustainable Development Wildlife and Fisheries Branch, 2017). In recent years, there has been growing concern over the state of the walleye fishery in Manitoba and its potential collapse, due to a number of factors including invasive species, eutrophication, fishing pressures, and climate change (Lake Winnipeg Quota Review Task Force, 2011; Wassenaar and Rao, 2012). This has increased the need to gain further insight into the physiological status of walleye in Manitoba, which has largely been understudied.

One location of potential concern for walleye is in the Delta Marsh, the largest coastal wetland on Lake Manitoba, Manitoba, Canada. Thirty-one species of fish use the marsh for spawning, rearing, and feeding (Wrubleski *et al*., 2018). While the marsh is widely known as an important wildlife habitat, especially for waterfowl (Batt, 2000), its importance as fishery habitat is not well understood. During the months of May to July, the Delta Marsh becomes inundated with invasive common carp (*Cyprinus carpio*) that use the marsh as a spawning ground, which has had detrimental effects on the ecosystem as a whole (Hnatiuk, 2006; Parks, 2006; Badiou and Goldsborough, 2010; Hertam, 2010). To mediate the negative impacts of common carp on the Delta Marsh, removable exclusion screens are put in place in the spring (e.g., late April) prior to the common carp migration into the marsh, and are removed in the late summer (e.g., early August). The installation of these exclusion screens has been successful in the remediation efforts of the Delta Marsh ecosystem (Wrubleski *et al*., unpublished data), but comes at a potential cost for large-bodied native fishes (>70 mm maximum body width) that are confined within the marsh during the warmest months of the year (e.g., Suthers and Gee, 1986). The Delta Marsh consists of a matrix of small and large-bodied bays, isolated ponds, and channels that range from less than 1 to 3 m in depth with an average depth of 1 m (Batt, 2000; Wrubleski *et al*., 2018). Thus, without access to the nearby cooler lake conditions, large native fishes held within the Delta Marsh may experience a sustained or repeated exposure to elevated water temperatures and low oxygen conditions for several months. Adult walleye growth is predicted to be optimal at temperatures between 20 and 24°C, and they tend to avoid temperatures exceeding 24°C (McMahon *et al*., 1984). Additionally, adult walleye abundance is highest at dissolved oxygen levels that exceed 3 to 5 mg l^-1^, and they can tolerate dissolved oxygen levels of 2 mg l^-1^ for only short periods of time (McMahon *et al*., 1984). Thus, physiological consequences from extended holding in the Delta Marsh during potentially high temperature and low oxygen periods could result in population-level effects for fish such as walleye.

The aim of the present study was to evaluate the physiological status of wild-caught walleye in the Delta Marsh using molecular tools. To this end, an annotated reference transcriptome for walleye gill tissue was produced using RNA-seq technology, providing the capacity to develop a suite of quantitative real-time PCR (qPCR) primers to measure the mRNA levels of several target genes to assess the physiological status of walleye. Wild-caught walleye confined in the Delta Marsh for approximately 1.5 and 3.5 months, constituting relatively short- and long-term holding in the marsh, were collected using a hoop net and sampled non-lethally for gill tissue in the first week of June and August of 2017. Over this sampling period, environmental conditions, including water temperature and dissolved oxygen, were recorded at the three primary corridors that connect the Delta Marsh to Lake Manitoba. Because walleye held under these conditions were expected to experience elevated water temperatures and low oxygen conditions, the transcriptomic patterns of 32 genes with roles in the temperature and general stress response, oxidative stress response, cell remodeling and repair, DNA repair, metabolism, hypoxia, and ion and acid-base regulation were measured. Overall, the molecular tools developed in the present study could be used to assess the physiological status of walleye in other Manitoba systems of concern, such as Lake Winnipeg, or more broadly, throughout their distribution (e.g., the Great Lakes of North America).

## Materials and Methods

### Delta Marsh sampling

In 2017, the exclusion screens in the Delta Marsh were lowered on April 25 and then lifted August 4. Walleye (*n* = 21) were sampled from Waterhen Creek (50.2319°N, 98.1154°W; Fig. 2) on the east side of the Delta Marsh on two occasions during this period, June 6–8, 2017 (*n* = 13) and August 1–4, 2017 (*n* = 8). Fish were captured in a 1.22 m diameter hoop net mounted on the exclusion structure at Waterhen Creek (Fig. 2). Briefly, a hoop net was positioned on the lake side of the exclusion structure and an exclusion screen was lifted temporarily to allow large fish from the marsh to pass through the exclusion structure into the net on the lake side. Bars within the exclusion screens were positioned 70 mm apart, but only fish with a width greater than 80 mm (range: 81–112 mm) were sampled, as they were most likely unable to pass through the screens and represent fish held in the marsh. Upon capture, fish were anesthetized using a 50 mg l^-1^ solution of tricane methanesulfonate (MS-222) buffered with sodium bicarbonate. Gill tissue was sampled by collecting the tips of 3–5 filaments from the 2^nd^ or 3^rd^ gill arch, that were placed into RNA*later* solution (Thermo Fisher Scientific, Waltham, MA, USA), and stored at 4°C overnight prior to storage at −80°C. Fish were implanted with an anchor tag to monitor any potential recaptures and then released back into the marsh.

**Figure 2.**
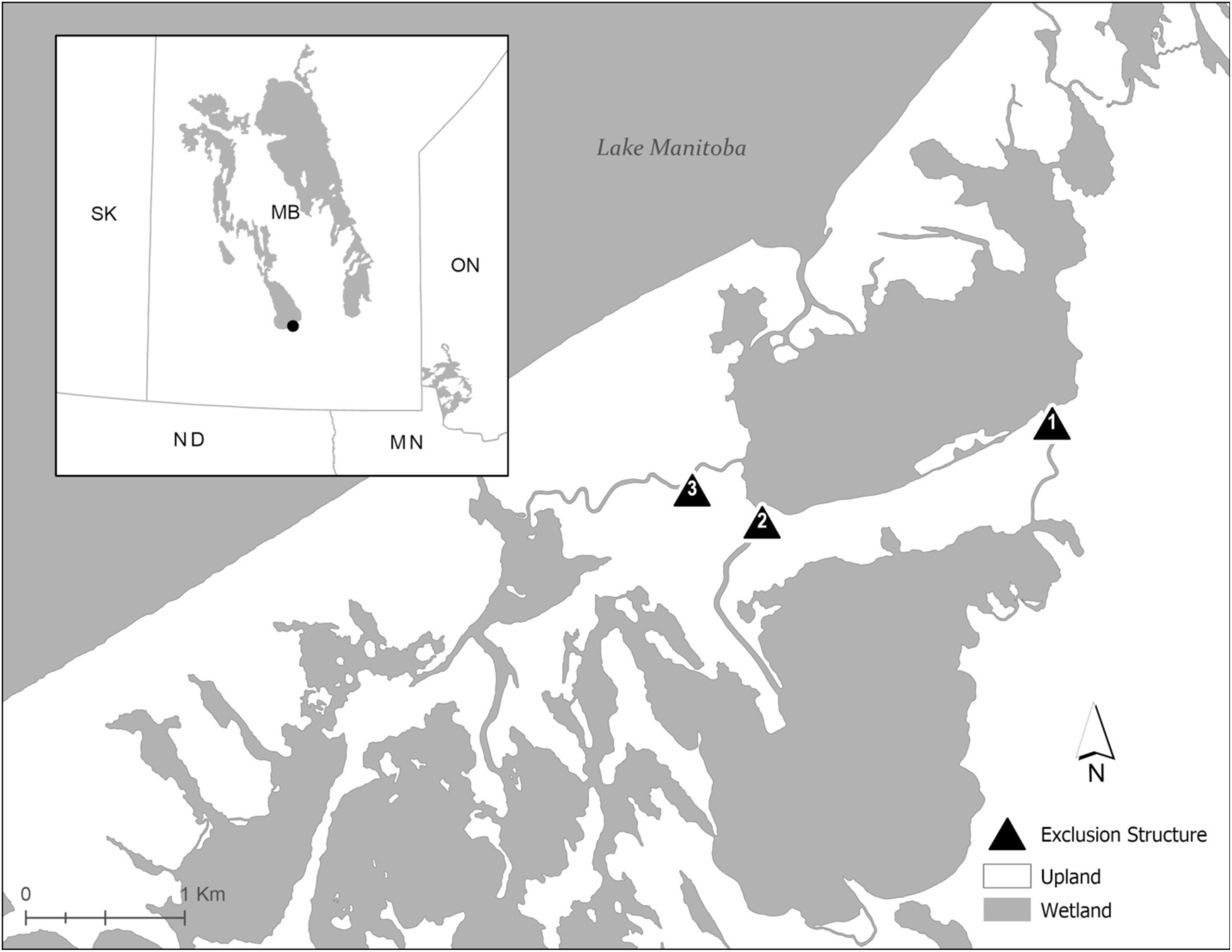
Map of the Delta Marsh region in Manitoba, Canada. Locations for water dissolved oxygen and temperature measurements are indicated by black triangles and include Fish Creek (1), Waterhen Creek (2), and Crooked Creek (3). Walleye (*Sander vitreus*) were non-lethally sampled in June and August from Waterhen Creek. The black circle on the inset map indicates the location of the Delta Marsh at the base of Lake Manitoba.

Dissolved oxygen (DO) and water temperatures were continuously monitored (hourly) beginning on May 12 until the end of the sampling using submerged HOBO Water Temperature and Dissolved Oxygen Data Loggers (UA-002-08 and U26-001, Onset, Bourne, MA, USA) at three locations: Waterhen Creek, Crooked Creek, and Fish Creek (Fig. 2). Each of these locations represent inflows into the Delta Marsh, as well as the location where fish were sampled (i.e., Waterhen Creek). Waterhen Creek constitutes one of the deepest areas within the marsh and carries upwards of 73% of the flow between the marsh and Lake Manitoba, while Crooked Creek and Fish Creek represent 11% and 3% of the flux, respectively (Aminian, 2015). Water oxygen saturation was calculated as: percent saturation = (100 × DO mg l^-1^)/C_p_, where C_p_ is equilibrium oxygen at nonstandard pressure (Mortimer, 1956). The cumulative number of days where conditions reached above 20 and 24°C (the lower and upper preferred temperature range for walleye), as well as below 30% oxygen saturation (i.e., hypoxic levels) were calculated for walleye sampled in June and August. It should be noted that dissolved oxygen values for the period of June 15–26, 2017 at Crooked Creek were continually at or below zero and were thus removed as they may represent a malfunction in the logger rather than true values.

### Lake Winnipeg sampling

Additional walleye were sampled across five locations in the Lake Winnipeg Basin, Manitoba, Canada, to include in the reference transcriptome (*n* = 10 total; *n* = 2 from each location): Dauphin River, Matheson, Riverton, Red River, Winnipeg River. Fish were collected by electrofishing at each location, with the exception of the Winnipeg River, where fish were angled by rod and reel. Walleye were sampled non-lethally for gill tissue as above. All fish collection and sampling techniques were in accordance with animal use protocols of Fisheries and Oceans Canada (FWI-ACC-2017-026) and the University of Manitoba (#F18-019).

### RNA extraction

Total RNA was extracted from the gill samples collected from both Lake Winnipeg walleye (*n* = 10) and Delta Marsh walleye (*n* = 21) using the RNeasy Plus Mini Kit (Qiagen, Toronto, ON, CA) following manufacturer’s instructions. The quantity and quality of RNA was determined using a NanoDrop One (Thermo Fisher Scientific) and by visualization on a 1% agarose gel, respectively.

### Transcriptome analysis and annotation

Fish were collected from multiple locations from the two main basins in Manitoba, Lake Manitoba (i.e., Delta Marsh) and Lake Winnipeg, to provide a diversity of individuals to include in a reference transcriptome for walleye. Equal amounts of total RNA were pooled from two samples from each of the six sampling locations (see above). The construction and sequencing of the cDNA library, as well as transcriptome assembly and annotation were performed by the McGill University and Genome Québec Innovation Centre (http://gqinnovationcenter.com) and the Canadian Centre for Computational Genomics (http://www.computationalgenomics.ca/), respectively. Prior to stranded mRNA isolation library construction using NEBnext library kits (New England Biolabs, Evry, France), the quality of the RNA pool was assessed on a Bioanalyzer 2100 (Agilent; R.I.N. = 8.6). Sequencing was performed using an Illumina HiSeq 4000 on a single lane and produced more than 326.8 million 100 base pairs paired-end reads. Trinity (Grabherr *et al*., 2011; Haas *et al*., 2013) and Trinotate (Bryant *et al*., 2017) were used for transcriptome assembly and annotation, respectively. Raw reads are available through the National Center for Biotechnology Information Sequence Read Archive database with the accession number SRP150633.

### Quantitative PCR

For walleye sampled in June and August from the Delta Marsh, the QuantiTect Reverse Transcription Kit (Qiagen) was used to synthesize cDNA from 1 µg of total RNA following the manufacturer’s protocol, with the exception that reaction volumes were scaled to a final volume of 40 µl. The relative abundance of 32 target genes (Table 1) was determined by qPCR. Sequences from the annotated walleye reference transcriptome were used to develop primers for target genes as well as five reference genes using Primer Express v.3 (Applied Biosystems, Thermo Fisher Scientific) (Table 1). To optimize reaction compositions, standard curves were generated for each primer set using cDNA synthesized from the RNA pooled from six individuals. PowerUP SYBR Green Master Mix (Applied Biosystems, Thermo Fisher Scientific) and a QuantStudio 5 Real-Time PCR System (Thermo Fisher Scientific) were used to carry out qPCR according to manufacturer’s instructions, with the exception that reaction compositions were scaled to 12 µl reactions. Target gene mRNA levels were normalized to the five reference genes (Table 1) and were expressed relative to the mean of the fish sampled in June using the delta-delta C_t_ method. For each reference gene, there was no effect of sampling period (i.e., June or August) on the mRNA abundance.

**Table 1:**
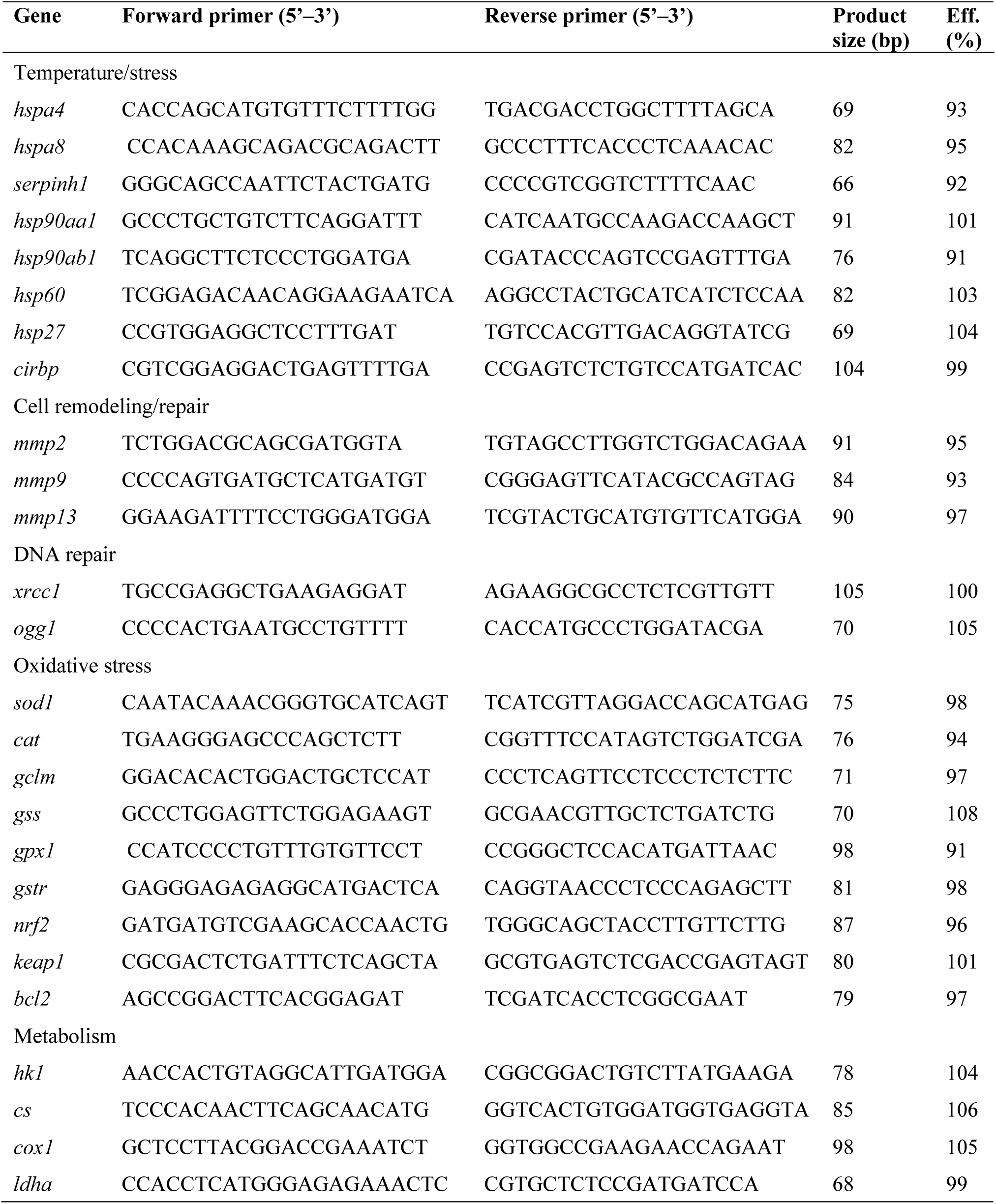
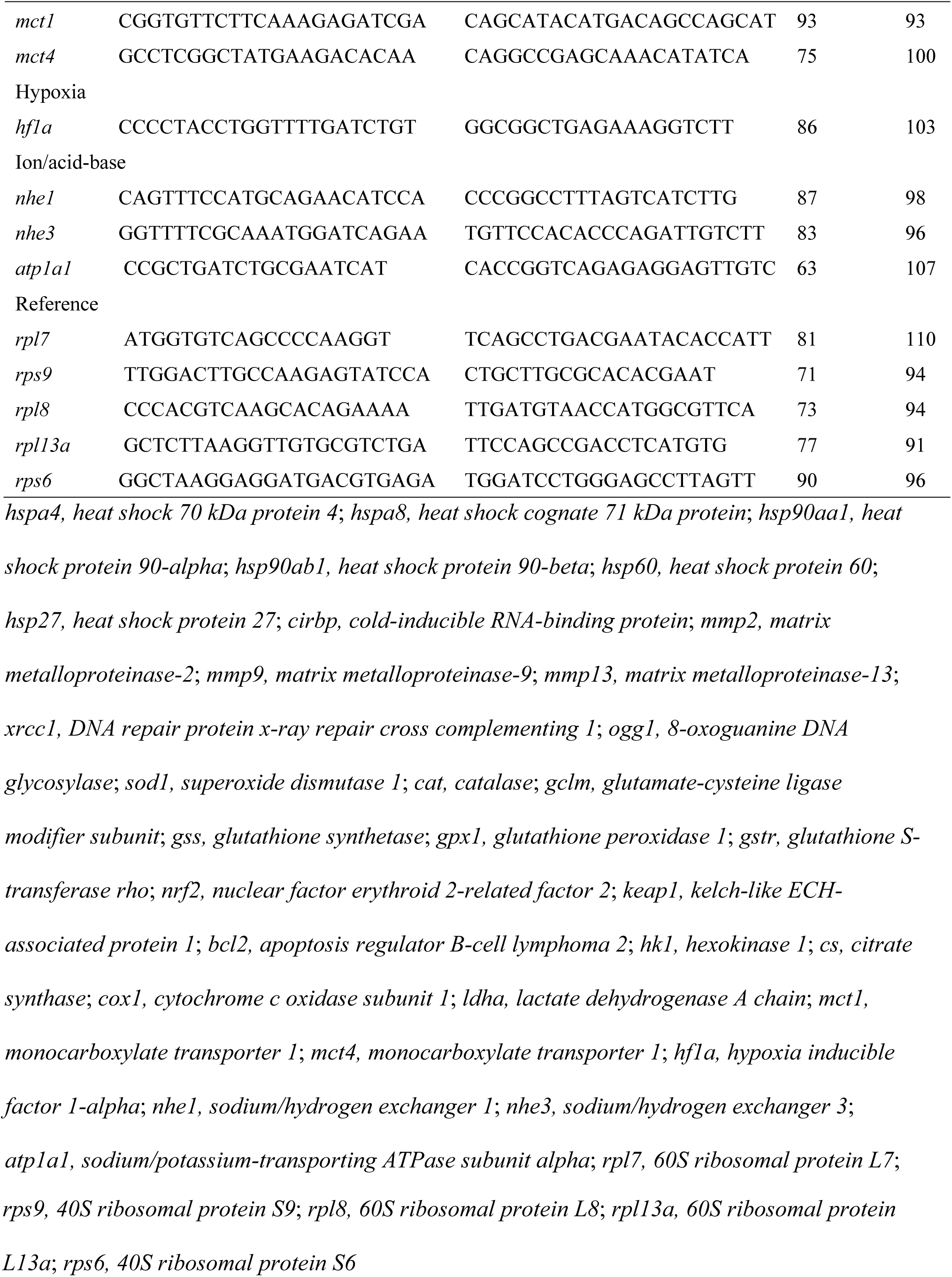
Oligonucleotide primers for quantitative PCR in walleye (*Sander vitreus*).

### Statistical analysis

All statistical analyses were run in R v3.6.1 (R Core Team, 2018), and the level of significance (α) was 0.05. Differences in the relative mRNA abundance of target genes between fish sampled in June and August were assessed by *t*-tests. The assumptions of equal variance and normality were tested for each variable using a Levene’s or Shapiro-Wilk’s test, respectively. If data were not equally variable across groups, a Welch’s Two Sample *t*-test was used. If data were not normally distributed, a Wilcoxon rank sum test was performed. In addition to individual *t*-tests, broad patterns of differential transcript expression among groups were analyzed by principal components analysis (PCA). A PCA was performed using the R package “FactoMineR” (Lê *et al*., 2008) and results were visualized using “factoextra” (Kassambara and Mundt, 2017).

## Results

### De novo assembled walleye reference transcriptome

The 326,849,617 reads from the pooled RNA-seq sample were assembled using Trinity. The assembly consisted of 438,162 transcript contigs that were clustered into 263,982 gene groupings with a median transcript length of 381 bp. In total, 198,990 transcripts and 86,539 genes were annotated using Trinotate.

### Water temperature and oxygen saturation

Similar water temperature profiles were measured across Waterhen Creek, Crooked Creek, and Fish Creek throughout the water sampling period of May 12 to August 4, 2017 (Fig. 3). Water temperatures began to rise in late May and early June and remained elevated (i.e., above ∼20°C) throughout the remaining water sampling period with the exception of late June when temperatures fell below 15°C. The maximum temperatures measured during the water sampling period were 26.7°C and 28.0°C on July 31 at Waterhen Creek and Fish Creek, respectively, and 27.4°C at Crooked Creek on July 28. Water temperatures reached above 24°C, the upper limit of the preferred temperature range for walleye (McMahon *et al*., 1984), on a cumulatively higher number of days for the August compared to June sampling periods, and at Crooked Creek and Fish Creek compared to Waterhen Creek (Table 3).

**Figure 3.**
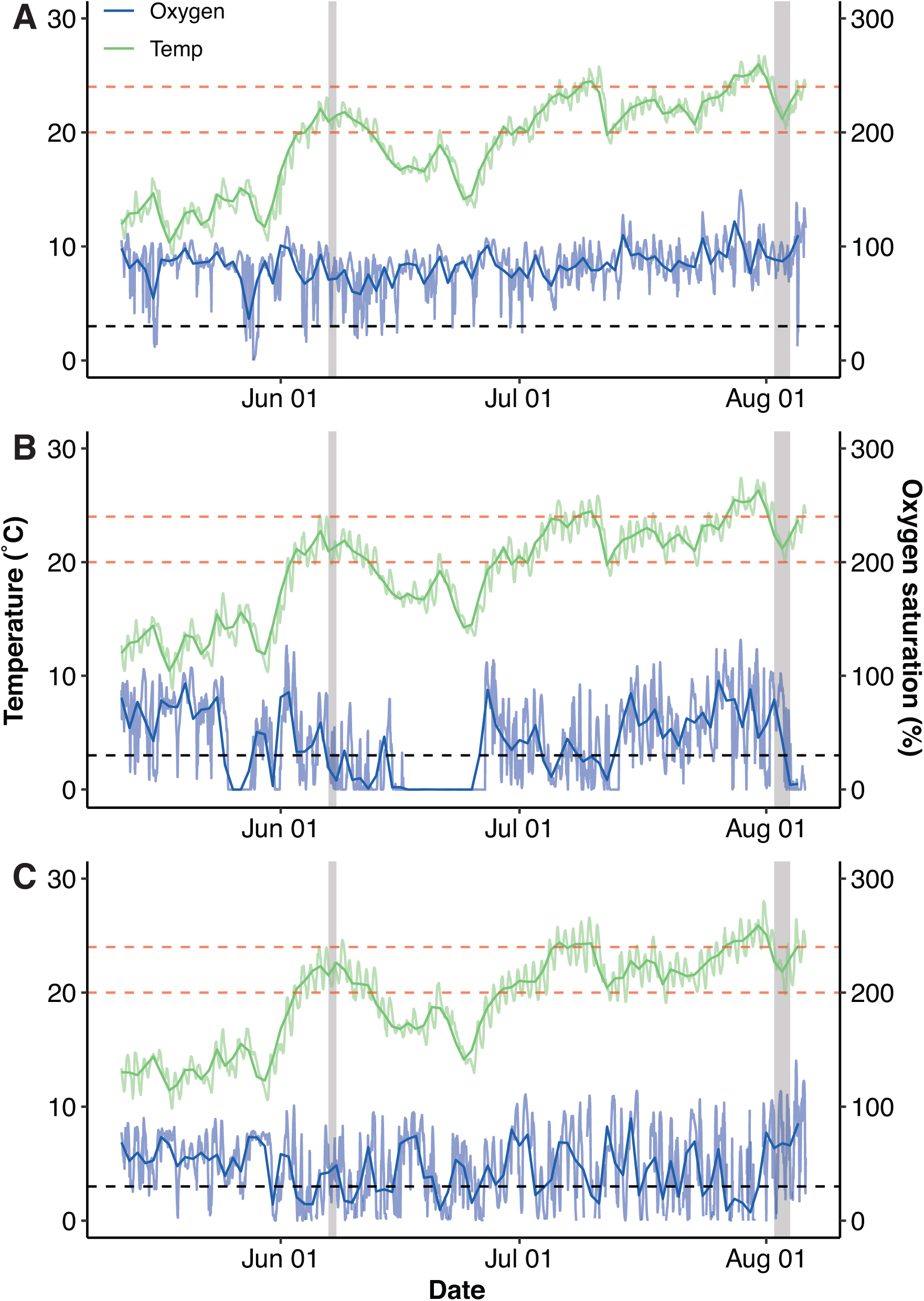
Water temperature and oxygen saturation at Waterhen Creek (A), Crooked Creek (B), and Fish Creek (C), in the Delta Marsh, Manitoba. Darker coloured lines represent daily means, and lighter coloured lines represent hourly measured values for temperature and oxygen saturation. The black hatched line represents 30% oxygen saturation, below which conditions would be considered hypoxic. The red hatched lines represent the upper and lower preferred temperature range (20–24°C) for walleye (*Sander vitreus*). Vertical grey boxes represent the sampling periods when walleye were collected.

**Table 2.**
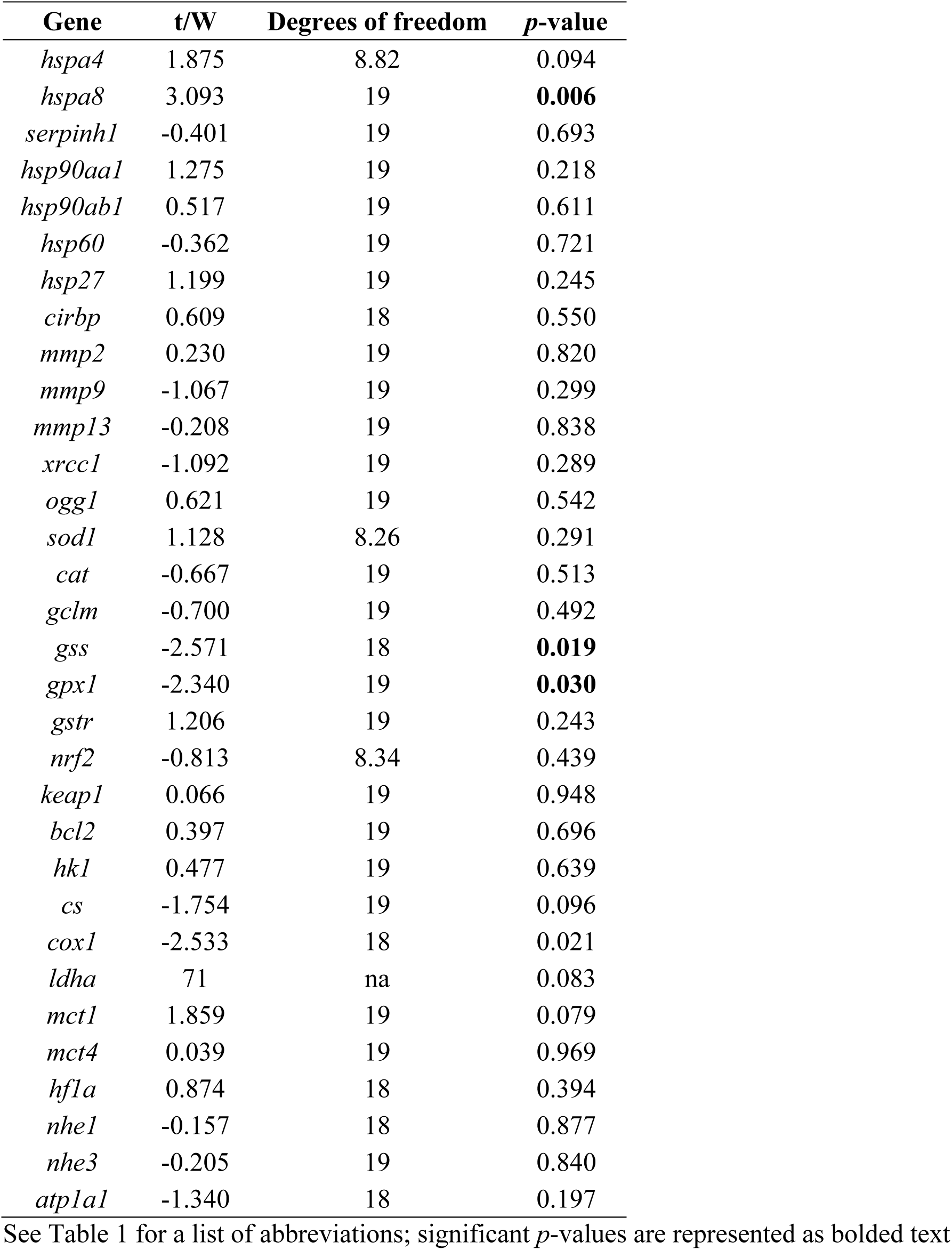
*t*-test results for the relative mRNA levels of 32 target genes from walleye (*Sander vitreus*) sampled in June and August from the Delta Marsh, Manitoba.

**Table 3.** Summary of water quality parameters for the periods prior to walleye (*Sander vitreus*) sampling in June and August 2017 for Waterhen, Crooked, and Fish Creeks. Water quality parameters are presented as cumulative number days above 20 or 24°C, and less than 30% O_2_ saturation for the entire period preceding fish sampling (i.e., June: May 12–June 8; August: May 12–August 4).

| Sampling period | Waterhen Creek |  | Crooked Creek |  | Fish Creek |  |
| --- | --- | --- | --- | --- | --- | --- |
|  | June | August | June | August | June | August |
| Days > 20°C | 7 | 52 | 8 | 54 | 7 | 52 |
| Days > 24°C | 0 | 15 | 1 | 18 | 2 | 23 |
| Days < 30% O<sub>2</sub> | 6 | 14 | 21 | 60<sup>1</sup> | 21 | 72 |

<sup>1</sup>Represents the minimum number of days < 30% O<sub>2</sub> saturation since values between June 15–26, 2017 were not available.

Water oxygen saturation levels varied across the three water sampling locations (Fig. 3). Water oxygen saturation levels reached below 30% for the fewest number of days at Waterhen Creek, compared to Fish and Crooked Creeks. Additionally, water oxygen saturation levels fell below 30% for a cumulatively higher number of days for the August compared to the June sampling periods across all three sites (Table 3).

### Transcriptional response to long-term holding within the Delta Marsh

Several mRNA transcripts of genes associated with heat stress and metabolism were differentially regulated in walleye held within the Delta Marsh for an extended period of time. Walleye sampled in August had significantly elevated levels of *heat shock cognate 71 kDa protein* (*hspa8*) mRNA (Fig. 4B; Table 2), and there was a trend for *heat shock 70 kDa protein 4* (*hspa4*) mRNA levels to also be elevated (*p* = 0.094; Fig. 4A), compared to fish sampled in June. The mRNA transcript abundance of genes associated with anaerobic metabolism, *lactate dehydrogenase A chain* (*ldha*) (Fig. 5A) and *monocarboxylate transporter 1* (*mct1*) (Fig. 5B) also tended to be elevated in walleye sampled in August compared to June (*p* = 0.082 and 0.078, respectively). Whereas the mRNA levels of genes associated with aerobic metabolism, *cytochrome c oxidase subunit 1* (*cox1*) (Fig. 5C; Table 2) and *citrate synthase* (*cs*) (Fig. 5D), were significantly, or tended to be (*p* = 0.096 for *cs*) lower in fish sampled in August compared to June. The mRNA transcript levels of genes associated with oxidative stress were elevated in walleye sampled during the earlier compared to later period following holding in the Delta Marsh. Both *glutathione synthetase* (*gss*) and *glutathione peroxidase 1* (*gpx1*) mRNA levels were significantly higher in walleye sampled in June compared to August (Fig. 6; Table 2).

**Figure 4.**
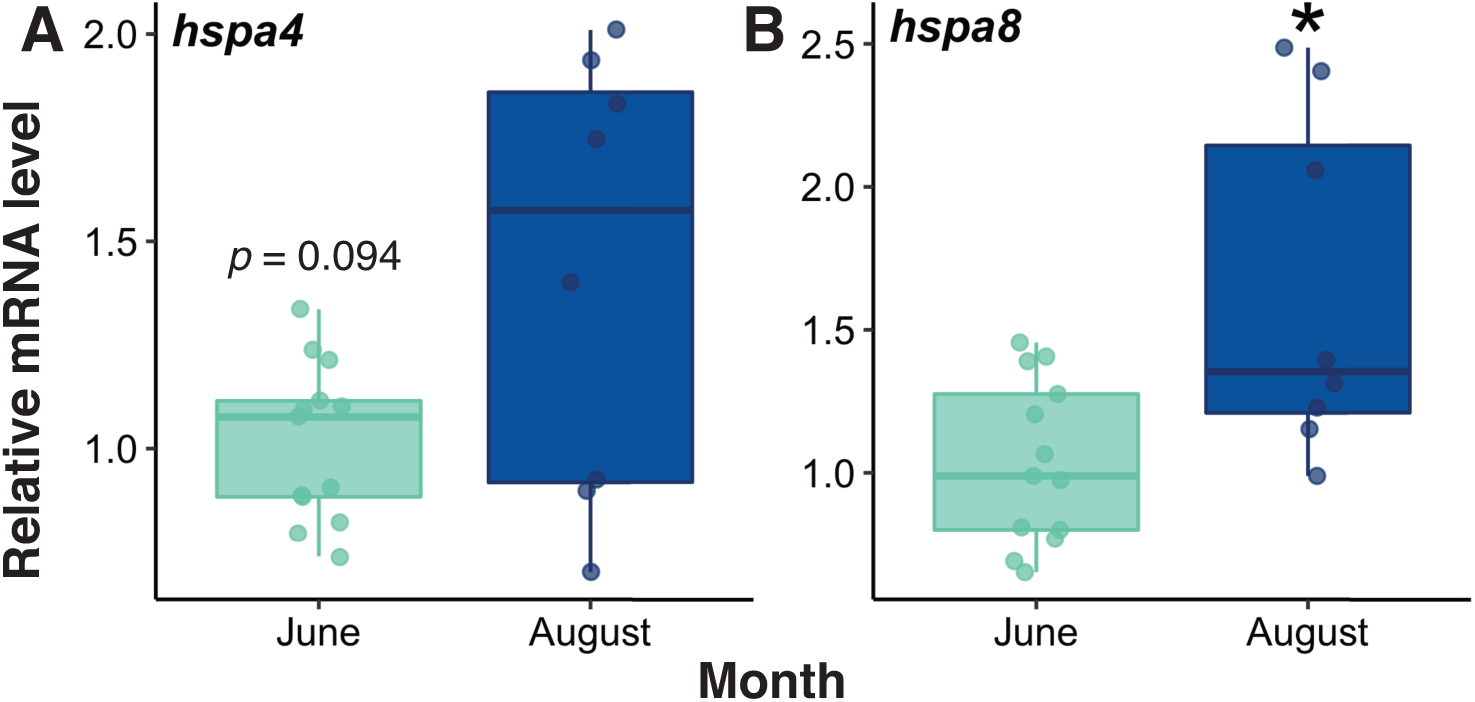
Relative mRNA levels for *heat shock 70 kDa protein 4* (*hspa4*; A), *heat shock cognate 71 kDa protein* (*hspa8*; B) of walleye (*Sander vitreus*) sampled from the Delta Marsh, Manitoba, in June (*n* = 13) and August (*n* = 8). An asterisk represents a significant difference between fish captured in June and August (*t*-test, see Table 2). Horizontal bars in the boxplot represent the median response value and the 75 and 25% quartiles. Whiskers represent ± 1.5 times the interquartile range, and each dot represents an individual response value.

**Figure 5.**
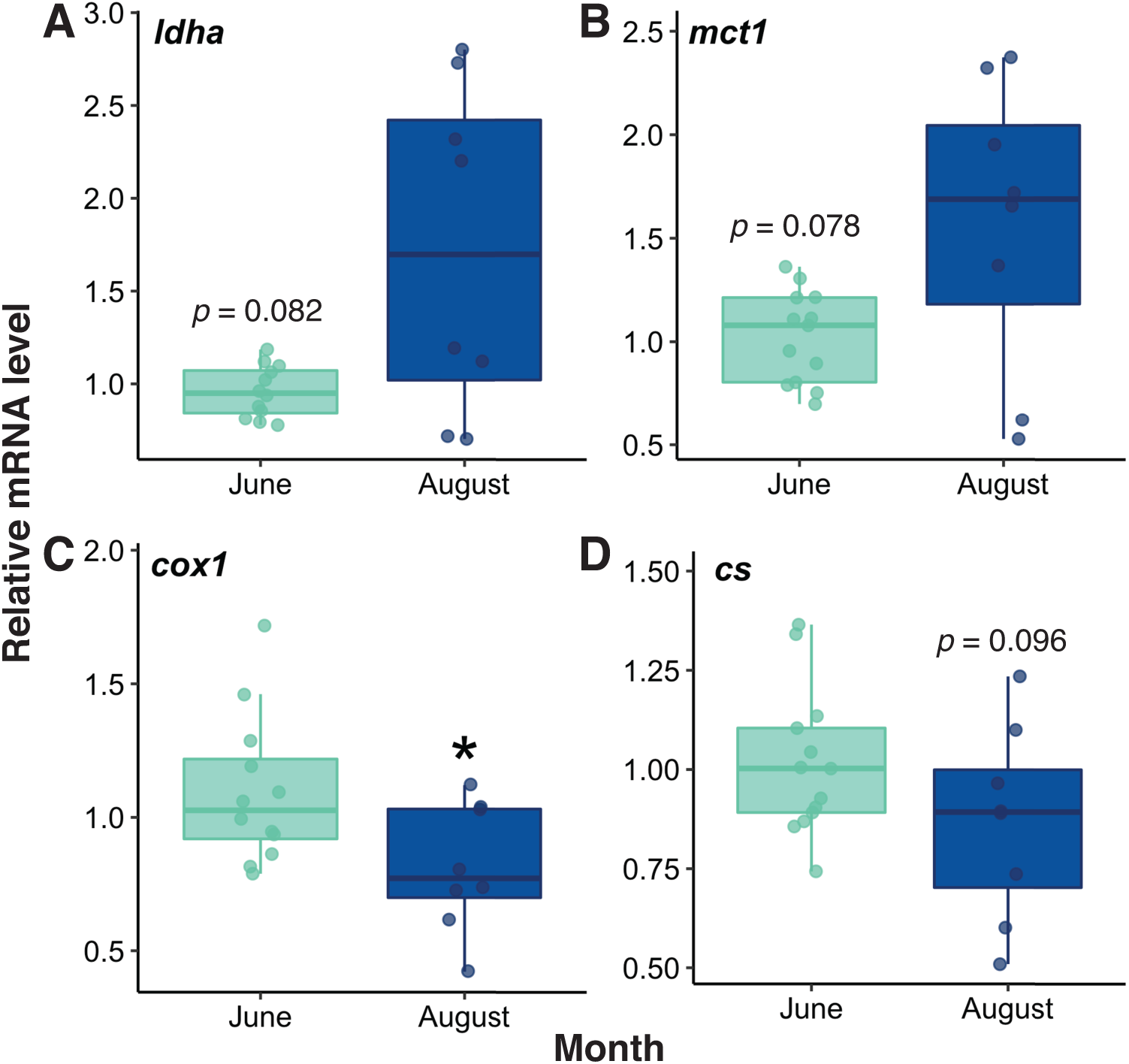
Relative mRNA levels for *lactate dehydrogenase A chain* (*ldha*; A), *monocarboxylate transporter 1* (*mct1*; B), *cytochrome c oxidase subunit 1* (*cox1*; C), and *citrate synthase* (*cs*; D) of walleye (*Sander vitreus*) sampled from the Delta Marsh, Manitoba, in June (*n* = 12–13) and August (*n* = 8). An asterisk represents a significant difference between fish captured in June and August (*t*-test, see Table 2). Horizontal bars in the boxplot represent the median response value and the 75 and 25% quartiles. Whiskers represent ± 1.5 times the interquartile range, and each dot represents an individual response value.

**Figure 6.**
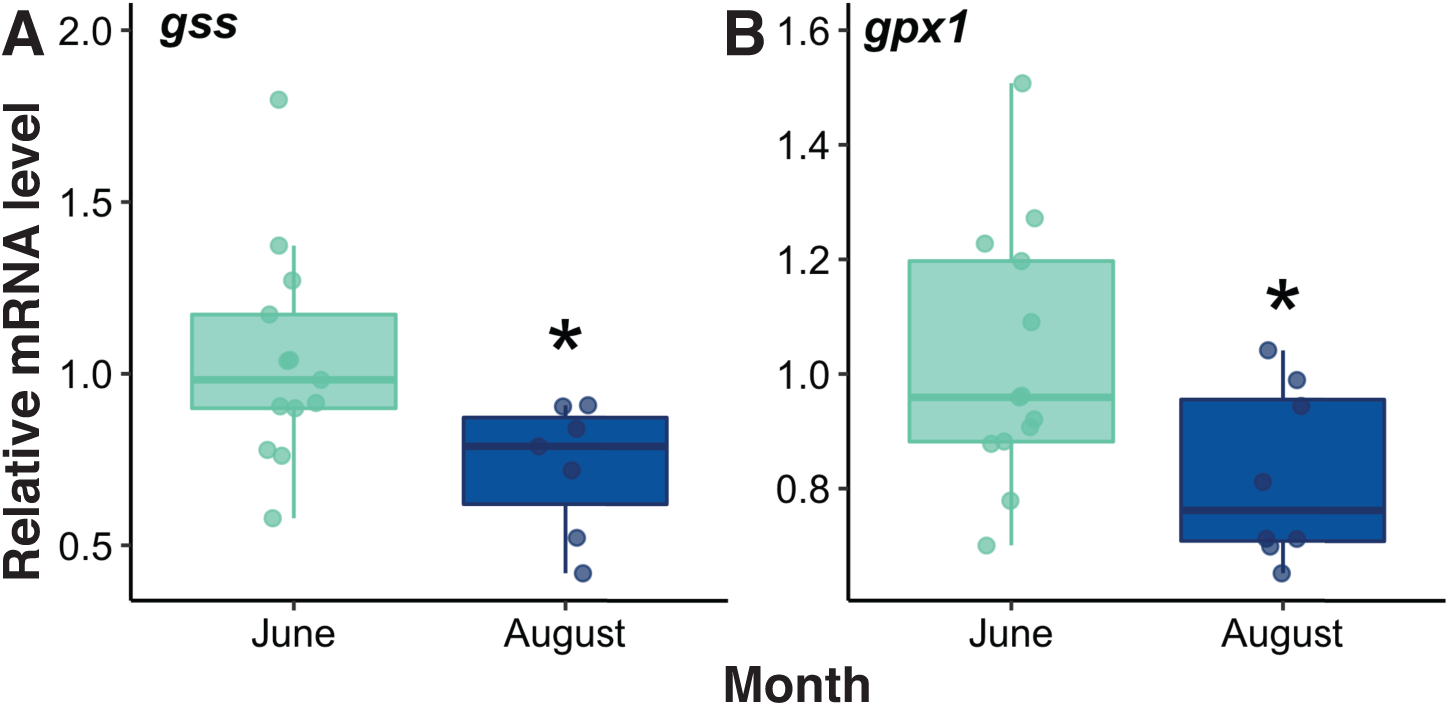
Relative mRNA levels for *glutathione synthetase* (*gss*; A), and *glutathione peroxidase 1* (*gpx1*; B) of walleye (*Sander vitreus*) sampled from the Delta Marsh, Manitoba, in June (*n* = 13) and August (*n* = 7–8). An asterisk represents a significant difference between fish captured in June and August (*t*-test, see Table 2). Horizontal bars in the boxplot represent the median response value and the 75 and 25% quartiles. Whiskers represent ± 1.5 times the interquartile range, and each dot represents an individual response value.

Using a PCA, clustering of walleye by sampling period occurred along the third principal component (PC), which explained 9.2% of the total variation, with fish sampled in August on the positive end and fish sampled in June at the negative end of the PC3 axis (Fig. 7A). Both PC1 and PC2 appeared to be related to interindividual variation in transcript levels. Of the 32 target genes examined, 16 genes contributed more to PC2 and PC3 than would be expected if all 32 genes contributed equally to these dimensions. Thus, only these 16 most highly contributing genes were represented as vectors along the second and third PCs for visualization purposes (Fig. 7B). The mRNA levels of genes associated with heat and general stress (*hspa8*, *hspa4, heat shock protein 90-beta* [*hsp90ab1*; Fig. S1A], *heat shock protein 27* [*hsp27*, Fig. S1B], and glycolysis and anaerobic metabolism (*hexokinase 1* [*hk1;* Fig. S1D], *ldha, mct1*) were positively loaded along PC3. Whereas the mRNA levels of genes associated with aerobic metabolism (*cox1*, *cs*) and oxidative stress (*gss*, *gpx1*, *glutamate-cysteine ligase modifier subunit* [*gclm*; Fig. S2B], *catalase* [*cat*; Fig. S2C]), as well as thermal stress (*heat shock protein 60* [*hsp60*; Fig. S2A]) were negatively loaded along PC3.

**Figure 7.**
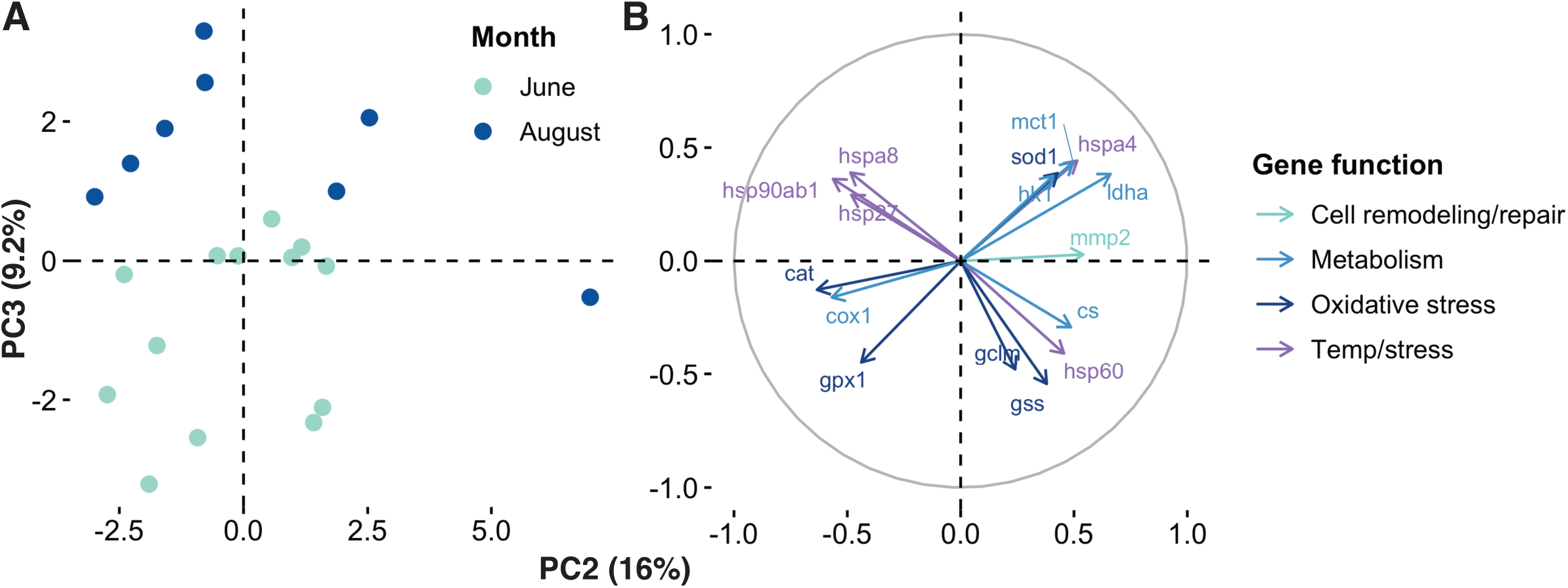
Principal components analysis (PCA) of the relative mRNA level for 32 target genes in walleye (*Sander vitreus*) sampled from the Delta Marsh, Manitoba, in June (*n* = 13) and August (*n* = 8). Clustering of walleye from June and August can be visualized along the 2^nd^ and 3^rd^ principal components (A). The top 16 highest contributing genes to PC1 and PC2 are coloured according to their biological function (B). See Table 1 for a list of gene name abbreviations.

## Discussion

In the present study, we successfully utilized a molecular approach to assess the physiological status of walleye in a location of conservation concern. Walleye held in the Delta Marsh for more than three months likely experienced increasing exposure to environmentally harsh conditions of high temperature and low oxygen. This was reflected in the transcriptional responses of fish held in the marsh until the end of the holding period (i.e., approximately 3.5 months for August sampled fish) when compared to fish sampled after a shorter period (i.e., approximately 1.5 months for June sampled fish). For example, a PCA revealed that long-term holding in the Delta Marsh resulted in regulation of the mRNA levels of temperature sensitive genes (e.g., *hspa8*, *hsp90ab1*, *hspa4*, *hsp27*), as well as a shift to decrease the transcription of genes associated with aerobic metabolism (e.g., *cox1*, *cs*) and increase the transcription of genes required for enhanced anaerobic glucose metabolism (e.g., *hk1*, *ldha*). Thus, holding walleye in the Delta Marsh during the period when the exclusion screens are in place could have sub-lethal consequences for large-bodied fish. This approach demonstrated the usefulness of developing molecular tools to examine the physiological status of a wild fish species in a conservation framework. Furthermore, the present study highlights the effectiveness of integrating a targeted gene-suite approach, facilitated by first producing an annotated transcriptome for the tissue of interest, with multivariate statistical analyses to lessen some of the potential confounding factors of handling and capture stress and high levels of interindividual variation in wild-fish species.

### Thermal stress response to holding in the Delta Marsh

Walleye exhibited a pronounced thermal response with long-term holding in the Delta Marsh, likely due to the increased temporal exposure to elevated water temperatures. Over the monitored period of May 12 to August 4, 2017 (the final day of fish sampling), water temperatures rose to above 24°C on 15–23 d, depending on the water sampling site. These conditions exceed the preferred (optimal) temperature range for walleye of 20 to 24°C (McMahon *et al*., 1984). Accordingly, the PCA indicated that of the eight thermally responsive genes examined, four HSPs representing different HSP families, *hsp90ab1*, *hspa8* (*hsc70*), *hspa4* (*hsph2*), and *hsp27* (*hspb1*), were positively loaded along PC3, suggesting higher levels of these transcripts in fish sampled in August compared to June. Both *hsp90ab1* and *hspa8* are constitutively expressed molecular chaperones involved in the refolding and stabilization of proteins and have been shown to be thermally responsive (Jeffries *et al*., 2014; Huang *et al*., 2018). The *hspa4* can act in synergy with *hspa8* either as a nucleotide exchange factor or as a chaperone itself (Dragovic *et al*., 2006; Mattoo *et al*., 2013). Categorized as a small HSP, *hsp27* may act as a primary line of defense under stress by neutralizing unfolded proteins and cooperating with other chaperones for further protein processing (Vos *et al*., 2008). Further, *hsp27* may help to stabilize cytoskeletal components such as actin, intermediate filaments, and microtubules (Landry and Huot, 1995; Vos *et al*., 2008), which could make it particularly important for maintaining gill structure during thermal stress. Similar increases in these HSPs have been found in other fish species exposed to thermal stress. Two species of adult Pacific salmon (sockeye salmon, *Oncorhynchus nerka*; pink salmon, *O. gorbuscha*) exhibited increased *hsp90ab1* mRNA levels in the gill following 5–7 d of holding at 19°C compared 13–14°C (Jeffries *et al*., 2012; Jeffries *et al*., 2014). Rainbow trout (*O. mykiss*) exposed to a moderate heat stressor (1°C increase day^-1^ from 18 to 24°C) also showed a significant elevation of *hsp90ab1, hspa8*, and *hspa4* mRNA abundance in the head kidney and/or liver (Li *et al*., 2017; Huang *et al*., 2018). Similarly, hybrid catfish (*Ictalurus punctatus* × *I. furcatus*) exposed to up to 102 h at 36°C exhibited elevated levels of *hsp90ab1* mRNA in the gill and liver, and elevations in *hspa8* and *hsp27* mRNA in the gill compared to fish held at 24°C (Liu *et al*., 2013). Together, the results of the present study suggest a sustained elevation in the heat stress response of walleye held in the Delta Marsh for an extended period of time and potentially indicate a lack of acclimation to the high thermal conditions.

Conditions within the Delta Marsh were relatively stable until May 29, 2017, when mean daily water temperatures began to rise. Mean daily water temperatures in the channels rose consistently across the three water sampling sites from approximately 12 to 22°C in a period of 8 d prior to the onset of fish sampling in June. A limited thermal response was observed in the mRNA levels of *hsp60* (*hspd1*), which was negatively loaded along PC3, suggesting higher transcript levels in fish sampled in June compared to August. Localized inside the mitochondria (Cappello *et al*., 2015), *hsp60* has been found to be thermally responsive in fishes. For example, *hsp60* mRNA levels were elevated in the head kidney and liver of rainbow trout exposed to a 6°C increase in temperature over 6 d (Li *et al*., 2017; Huang *et al*., 2018), as well as in the liver of catfish held at 36°C for up to 5 d compared to 24°C (Liu *et al*., 2013). Similarly, *hsp60* mRNA levels increased initially but decreased over time following a heat challenge in snow trout (*Schizothorax richardsonii*), suggesting a role for *hsp60* in the short-term response to thermal stress (Barat *et al*., 2016). Because fish sampled in June were not compared to a group prior to the 10°C increase in temperature, a broader thermal response may have occurred at this time that would not have been captured by the present study. Nevertheless, the higher levels of *hsp60* mRNA in fish sampled in June compared to August suggest that *hsp60* may play an important role in the initial response to thermal stress.

### Oxidative stress response to holding in the Delta Marsh

The increase in temperature preceding the June sampling period may have contributed to an oxidative stress response. Thermal stress can lead to the excessive production of reactive oxygen species (ROS), leading to oxidative damage in fish (An *et al*., 2010; Kim *et al*., 2014). In response, the antioxidant defense system includes enzymes to scavenge ROS to prevent damaging oxidative stress. Both *gss* and *gpx1* mRNA levels were significantly higher in fish sampled in June compared to August, and together with *cat* and *gclm*, were negatively loaded along PC3 suggesting elevated mRNA abundance of these genes in fish sampled in June compared to August. Similar increases in antioxidant enzymes have been observed in other fishes exposed to thermal stressors. For instance, increases in the mRNA levels of *gpx1* and *cat* were observed in the liver of black porgy (*Acanthopagrus schlegeli*) exposed to an increase of 10°C at a rate of 1°C day^-1^ (An *et al*., 2010). Interestingly, the liver mRNA levels of *gpx1* were lower in Atlantic salmon (*Salmo salar*) held at 17 and 19°C compared to 13 and 15 °C for 45 d (Olsvik *et al*., 2013), suggesting that *gpx1* mRNA levels may have been suppressed in walleye sampled in August, rather than elevated in fish sampled in June. Similarly, in an Antarctic fish, the bald notothen (*Pagothenia borchgrevinki*), liver *cat* mRNA abundance was elevated in response to an acute (12 h) increase in temperature, but returned to basal levels following a long-term (3 wk) exposure to elevated temperatures (Carney Almroth *et al*., 2015). Of the antioxidant genes investigated in this study, only *superoxide dismutase 1* (*sod1*) mRNA levels tended to be elevated in fish sampled in August following long-term holding in the Delta Marsh (i.e., was positively loaded along PC3; Fig. S1C). Together, the results of the present study, as well as those of previous studies, suggest that the antioxidant response to elevated temperature may be transient, with decreased effectiveness during long-term exposure to thermally harsh conditions. To test this hypothesis, future studies should assess whether this transient antioxidant response is consistent across other walleye tissues (e.g., liver), and whether chronic exposure to elevated temperatures results in a concomitant increase in ROS and oxidative damage.

### Metabolic response to holding in the Delta Marsh

In addition to increases in temperature in the Delta Marsh, walleye likely also experienced low oxygen conditions, that at times reached hypoxic or anoxic levels in certain areas of the marsh. For instance, Waterhen Creek reached below 30% oxygen saturation (i.e., hypoxic levels) on 6 and 14 d for fish sampled in June and August, respectively, and likely represents one of the ‘best case scenario’ for walleye in the Delta Marsh as it is one of the deepest areas in the marsh and also carries the greatest flows between Lake Manitoba and the marsh (Aminian, 2015). At shallower sites, such as Crooked Creek and Fish Creek, oxygen levels fell below 30% saturation on 21 and 60–72 d for the periods prior to June and August samplings, respectively. Consequently, walleye metabolism was likely regulated to match low oxygen conditions experienced with longer holding in the marsh. Indeed, walleye sampled in August compared to June tended to have elevated mRNA levels of genes that may be associated with anaerobic glycolysis, including *ldha*, *mct1*, and *hk1*, which were positively loaded along PC3. In contrast, the mRNA levels of *cox1* were significantly lower in fish sampled in August compared to June, and together with *cs*, were negatively loaded along PC3, suggesting that genes associated with aerobic metabolism were reduced in fish sampled in August compared to June. This shift from aerobic to anaerobic metabolism at a molecular level (i.e., mRNA) during exposure to hypoxic conditions is a typical response in fishes (Richards, 2009). Similar increases in *ldha* and *mct1* mRNA in the gill were observed in zebrafish during short-term exposure to hypoxia (48 or 96 h) (Ngan and Wang, 2009). Adult zebrafish also exhibited elevations in the mRNA levels of genes associated with glycolysis and a reduction in the mRNA levels of genes involved in the Krebs cycle and the electron transport chain in response to 3 wk of holding in severe hypoxic conditions (van der Meer *et al*., 2005). Together, these results suggest that walleye held in the Delta Marsh may have been exposed to increasing periods of low oxygen availability, resulting in a shift in the regulation of metabolic pathways favoring anaerobic over aerobic mechanisms.

### Considerations for study design within a conservation physiology framework

The sampling of wild fish to examine the impacts of environmental and anthropogenic stressors provides valuable information regarding the physiological responses of fish in their natural setting, particularly for conservation purposes. However, in comparison to controlled laboratory studies, field studies introduce a level of variation that could potentially mask certain physiological responses of interest. For instance, variation in the physiological responses of fish can be affected by capture techniques and differences in the environmental conditions experienced prior to capture, as well as factors such as fish sex and age, that may not be able to be easily controlled or accounted for in the field (Simmons *et al*., 2015). One method to account for this interindividual variation in the physiological responses of wild fish is by increasing the sample size to increase statistical power (Simmons *et al*., 2015; Oomen and Hutchings, 2017). Unfortunately, increasing the sample size is not always possible in systems where access to fish is limited to certain periods of the year or the area is sparsely populated (e.g., Jeffrey *et al*., 2019). We suggest an alternative or complementary approach for molecular studies, in which a gene-suite approach is paired with multivariate statistics. In the present study, a high level of interindividual variation was evident for each individual gene as well as in the first two principal components (i.e., group clustering was not evident until the third PC). This made it challenging to detect statistically and biologically meaningful differences in the mRNA levels of genes of interest, even with a reasonably large sample size (i.e., *n* = 8–13). By selecting a total of 32 target genes, with multiple genes representing a biological response (e.g., oxidative stress, metabolism, heat stress response), and utilizing a multivariate statistical approach, we were able to discern different patterns of regulation in the physiological responses among our sampling groups (i.e., June *vs*. August sampled fish) along the third PC. Additionally, we attempted to target genes of interest that would be less sensitive to acute handling or capture stressors and thus would be more indicative of the conditions experienced prior to capture. Together, we believe that examining the mRNA levels of a suite of genes in combination with multivariate statistics provides a useful and informative approach for field studies, and in particular, within a conservation physiology framework.

### Conclusion

Overall, the results of the present study suggest that extended holding of large-bodied walleye in the Delta Marsh could have sub-lethal consequences due to their increasing exposure to harsh temperature and oxygen conditions. The present study identified a number of molecular biomarkers in non-lethally sampled wild fish that were sensitive to elevated levels of temperature and low oxygen under both short-term (i.e., *hsp60*, *gss*, *gpx1*) and long-term (i.e., *hspa8*, *hsp90ab1*, *cox1*) holding conditions. The results of the present study highlight the usefulness of developing molecular tools, specifically using transcriptomic techniques, to examine the physiological status of wild fish. By establishing a reference transcriptome for walleye, a number of possible biomarkers of stress could be selected across several physiological systems to allow for broad patterns of responses to be observed. Additionally, a gene-suite approach in combination with multivariate statistics may lessen the influence of potential confounding factors due to field capture techniques. Together, the results of the present study provide valuable knowledge regarding the physiological status of large-bodied walleye held in the Delta Marsh when the common carp exclusion screens are in place. These results could inform conservation decisions by managers, and more broadly, provides additional evidence for the usefulness of molecular techniques in conservation biology.

## Acknowledgements

We would like to thank several individuals for their help with fish sampling including J. Sutherby, W. Bugg, C. Wong, T. Naaykens, K. Clarete, C. Wlasichuk, and A. Caskenette. Thanks also to M. Gaudry for his help with RNA extractions, and to S. Witherly for producing the map of the Delta Marsh. Finally, we thank Dr. E. Gonzalez from the Canadian Centre for Computational Genomics who carried out the bioinformatics work.

