## Supplementary Materials for "Applying a gene-suite approach to examine the physiological status of wild-caught walleye (*Sander vitreus*)"

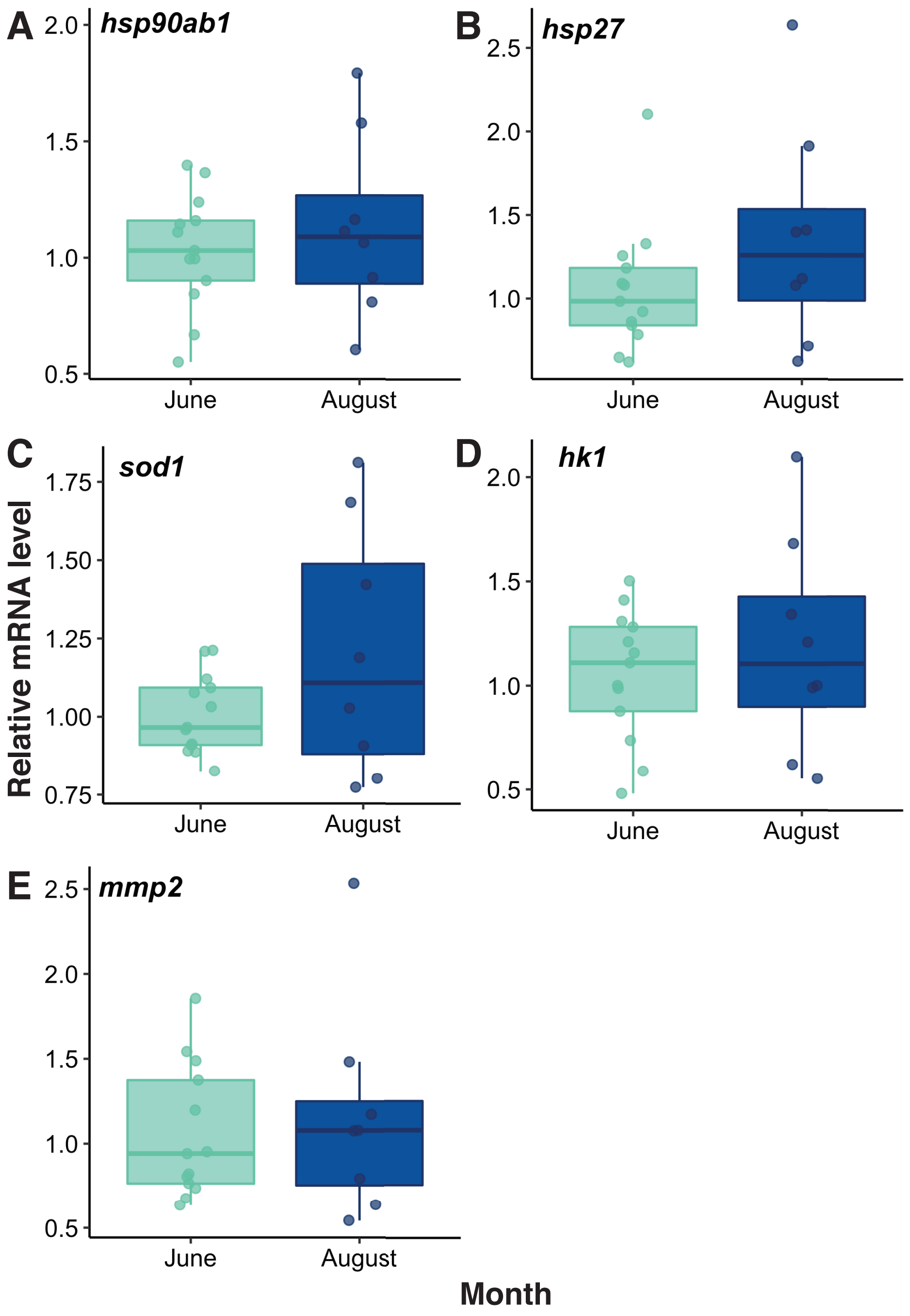


Figure S1. Relative mRNA levels for *heat shock protein 90-beta* (*hsp90ab1*; A), *heat shock protein 27* (*hsp27*; B), *superoxide dismutase 1* (*sod1*; C), *hexokinase 1* (*hk1*; D), and *matrix metalloproteinase-2* (*mmp2*; E) of walleye (*Sander vitreus*) sampled from the Delta Marsh, Manitoba, in June ($n$ = 13) and August ($n$ = 8). Horizontal bars in the boxplot represent the median response value and the 75 and 25% quartiles. Whiskers represent ± 1.5 times the interquartile range, and each dot represents an individual response value.

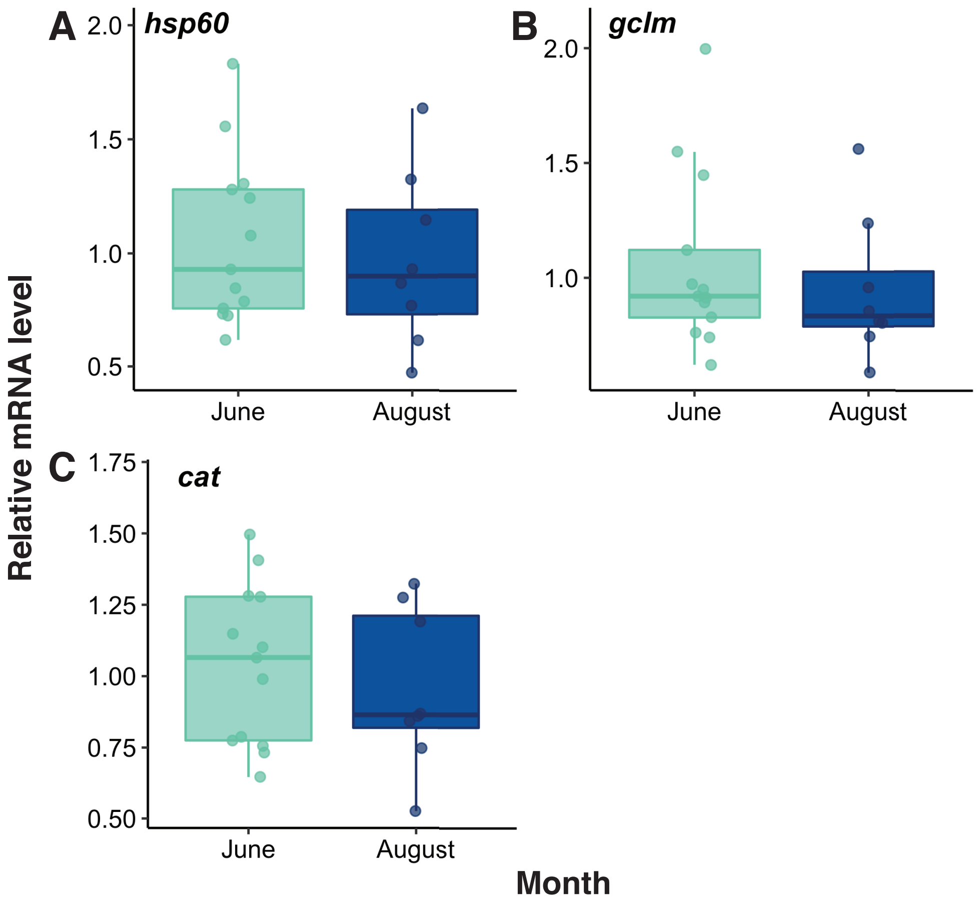


Figure S2. Relative mRNA levels for *heat shock protein 60* (*hsp60*; A), *glutamate-cysteine ligase modifier subunit* (*gclm*; B), and *catalase* (*cat*; C) of walleye (*Sander vitreus*) sampled from the Delta Marsh, Manitoba, in June ($n$ = 13) and August ($n$ = 8). Horizontal bars in the boxplot represent the median response value and the 75 and 25% quartiles. Whiskers represent ± 1.5 times the interquartile range, and each dot represents an individual response value.
